# A solvable model of entrainment ranges for the circadian rhythm under light/dark cycles

**DOI:** 10.1101/683615

**Authors:** Ren Takakura, Masatoshi Ichikawa, Tokitaka Oyama

**Affiliations:** Department of Advanced Mathematical Sciences, Graduate School of Informatics, Kyoto University, Yoshida-honmachi, Sakyo-ku, Kyoto 606-8502, Japan; Department of Physics, Graduate School of Science, Kyoto University, Kitashirakawa-oiwake-cho, Sakyo-ku, Kyoto 606-8502, Japan; Department of Botany, Graduate School of Science, Kyoto University, Kitashirakawa-oiwake-cho, Sakyo-ku, Kyoto 606-8502, Japan

## Abstract

A continuous model for entrainment of the circadian clock responding to light/dark cycles is analytically studied. The circadian rhythm is entrained to light/dark cycles in a certain range of periods including the 24 hour period. Entrainment ranges vary among organisms; plants show much larger ranges than mammals. To analyze entrainment manners, we exploit a simplified model in which the angular velocity of the circadian rhythm is modulated by a sinusoidal function of phase difference between the circadian rhythm and reference phases. This model contains only one parameter called entrainment strength (*K*). For light and dark states, we set two reference phases. Using this model, we characterized the entrainment manners of circadian rhythms under non-24 hour light/dark cycles. The conditions of the *K* value for the entrainment were analytically calculated and a phase diagram showing entrainment/non-entrainment boundaries was drawn. In the diagram we found a region where disordered orbitals emerge. Then, critical values for the entrainment and the disordered irregular motion of the oscillation were derived. Simulation results near a critical value were comparable with the experimental results of the entrainment manners of plant circadian rhythms, suggesting the compatibility of self-oscillation and a strong light/dark response in plants. The diagram clearly represents an overview of the relationship between entrainment strengths and entrainment manners of circadian rhythms in general, and enables us to uniformly compare strengths of periodic stimuli in the environment and degrees of responsiveness for the stimuli among various circadian rhythms.

## Introduction

Most organisms including plants, animals, and even bacteria have circadian rhythms to adapt to their environments with day-night cycles. Circadian systems exhibit self-oscillatory properties with an ability to synchronize to the cyclic environments. Appropriate synchronization of circadian systems is essential for their functions. These biological oscillation systems have been studied from various aspects: molecular, physiological, and mathematical ones (1–5).

The modern mathematical treatment of synchronization phenomena was initiated in the studies by Arthur Winfree (6,7). Following these, Yoshiki Kuramoto built a theoretical milestone in the field (8,9). Various phase related phenomena converged and were connected in the world of statistical mechanics through the Kuramoto model, including reduction theory. Following this, theoretical works on synchronization phenomena have utilized the framework of the Kuramoto model for its strengths in generality and applicability. On the other hand, more concrete models have been demanded in order to deeply understand complex synchronization behaviors and to unpack biochemical reaction mechanisms. Entrainment (synchronization) phenomena of circadian rhythms have been mathematically analyzed through two fundamental tools: the discrete model and the continuous model (1). The discrete model is based on the phase-shift as a response to pulse-like stimuli. This model has been successfully applied to phase shifts of circadian rhythms induced by periodic pulses such as skeleton photoperiods. However, the natural light/dark cycles are complete photoperiods that are unlike the simple pulse-like stimuli of lights. The continuous model is based on the modulation of a free-running period (*FRP*). The idea of this model seems to be more suitable for the entrainment of circadian rhythms in light/dark conditions. To assess capabilities of entrainment to complete photoperiods, non-24 hour light/dark cycles (*T*-cycles) have been experimentally applied to various circadian phenomena of animals, plants, and cyanobacteria, and theoretical models to understand those entrainment manners have been demanded (10–15). The molecular bases of circadian behaviors under *T*-cycles were also studied (16–18). Although continuous models proposed for these entrainment phenomena were numerically complicated in quantitative evaluation, Ermentrout and Rinzel (1984) proposed a simple continuous model to quantitatively explain the entrainment phenomena of firefly flashing rhythms to external periodic flashing lights (19). This model can be regarded as a direct application of the Kuramoto model on the experimental results, i.e., the external force with continuous phase modulates the angular velocity of flashing rhythms.

Here we propose a solvable continuous model with two states of external forces to analyze the entrainment phenomena of circadian rhythms in light/dark conditions. Instead of supposing external forces with a continuous phase, we assume a switching of external forces dependent on the light/dark conditions. Through this model, we quantitatively evaluate the capabilities of entrainment of circadian rhythms to light/dark cycles with various period lengths (*T*-cycles). Then we show that the results of this model correspond to the experimental results of heterogeneous cellular circadian rhythms in the same plant.

## Model

The Ermentrout/Rinzel model (19) for the entrainment of flashing rhythms to light/dark cycles is described by the following equations,

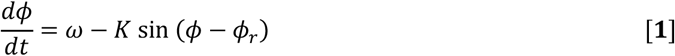

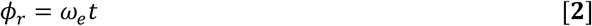

where *φ* is phase of flashing rhythms, *φ*_*r*_ is the reference phase of external force, *ω* is intrinsic angular velocity of flashing rhythms, *ω*_*e*_ is angular velocity of external force, and *K* is entrainment strength. The angular velocity can be expressed as

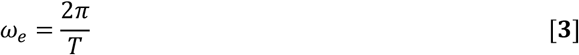

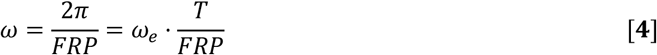

where *T* is the period length of light/dark cycles. In this study, *FRP* means a free-running period of a rhythm without any external forces (*K* = 0). When *φ*_*r*_ increases at a constant velocity, which means that the light/dark cycles give the continuous phase as temporal information to the rhythms, an entrainment orbit and the range of *K* for the entrainment were revealed by the ordinary differential equation (20). When applying this model to the entrainment of circadian oscillations under light/dark cycles (*φ* is phase of circadian oscillations, *ω* is intrinsic angular velocity of circadian oscillations), the assumption of “continuous phase information of the environmental cycles from 0 to *2π*” sounds to be unlikely. There are only two states: light and dark. As a simple assumption, we introduce two-state reference phases in our model.

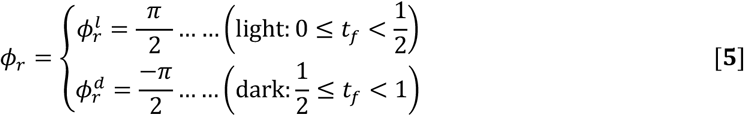

where 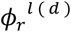 expresses the state of *φ*_*r*_ in light (dark), and (*t*_*f*_ expresses the fractional part of *t* (e.g. when *t* = 3.7 days, *t*_*f*_ = 0.7 days). In Eq. **3**, we normalize *ω*_*e*_ as *2π* and light/dark period lengths (*T*) as 1 day. Then, Eq. **4** is modified as

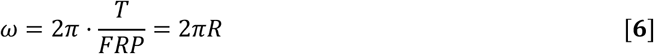

where *R* is the ratio of a light/dark period length to a free-running period. Eq. **1** is individually integrated under the assumption of Eq. **5** and roughly divided into two cases.

Case 1. 0 ≤ *K* ≤ *ω*

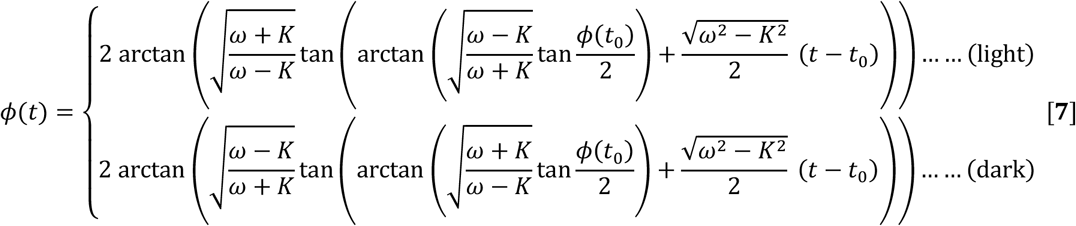

Case 2. *K* > *ω*

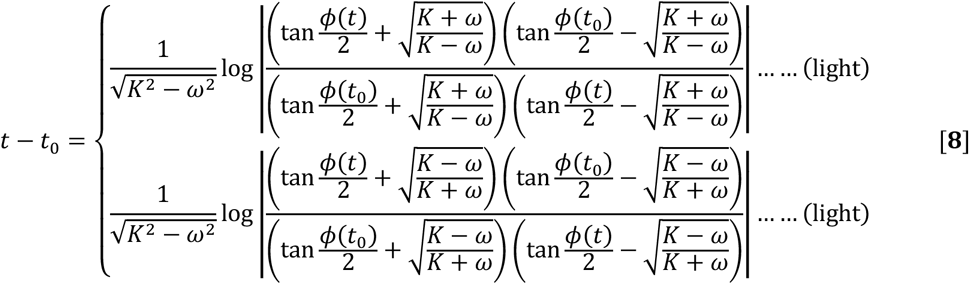

Strictly, *φ*(*t*) with *K* = ω is calculated by the limit value of Eq. **7**. See Appendix A for explicit description.

## Results and Discussion

We simulated various trajectories of *φ*(*t*) of circadian oscillations in light/dark cycles by changing the parameters (*K*, *R*) of our model. Then each trajectory is categorized into four types of oscillation characteristics: free-running, non-entrainment without phase jump, non-entrainment with phase jump, and entrainment (Fig. 1, phase jump means that 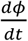 becomes negative and a kind of reverse motion of circadian oscillation occurs).

**Figure 1.**
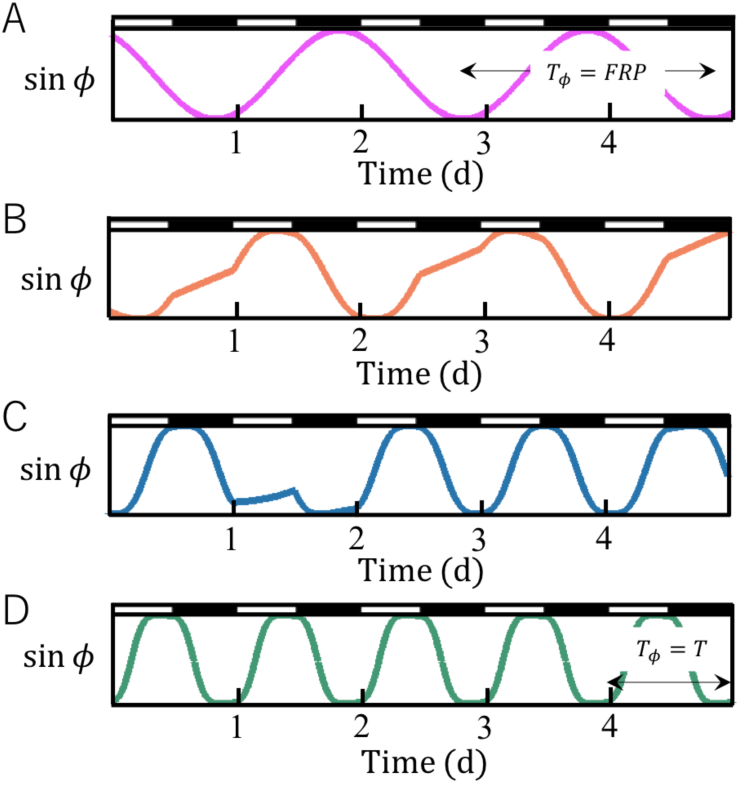
Four characteristic manners of oscillation. (A) Free-running (*K* = 0): Circadian oscillations are not affected by light/dark cycles and *T*_*φ*_ equals *FRP*. (B) Non-entrainment without phase jump (*K* < *K*_*c*_ and 0 ≤ *K* ≤ *ω*): Circadian oscillations are modulated by light/dark cycles. *T*_*φ*_ changes every cycle, but any *T*_*φ*_ does not equal *T*. The angular velocities of oscillations remain positive. (C) Non-entrainment with phase jump (*ω* < *K* < *K*_*c*_): Circadian oscillations are affected by light/dark cycles and *T*_*φ*_ does not equal *T*. Negative angular velocities of circadian oscillations, which can be regarded as “phase jump,” are observed. (D) Entrainment (*K* ≥ *K*_*c*_): Circadian oscillations are entrained to light/dark cycles and *T*_*φ*_ equals *T*. Open and filled rectangles denoted above the graph represent light and dark durations, respectively.

*K*_*c*_ means the critical value of *K* at which transition from “non-entrainment” to “entrainment” occurs. Behaviors of circadian oscillations are determined by (*K*, *R*), where 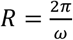 from Eq. **6**. Under any conditions of *R* the circadian oscillations are entrained to light/dark cycles when the *K* value is large enough. We analyzed relationships between (*K*, *R*) and oscillation behaviors, then we classified: into three types according to the manners of change in oscillation behaviors with increasing *K* (Figs. 2, 3). For the classification, we set two criteria: entrainment by increasing/decreasing of period lengths of circadian oscillation, and occurrence/non-occurrence of “non-entrainment with phase jump.” When the *FRP* of circadian oscillation is shorter than the periods of light/dark cycles (*T*), namely *R* > 1, the circadian oscillation can be entrained by increasing its period length. In this condition, we did not find phase jump of oscillation in any simulations. When *R* < 1, the circadian oscillation can be entrained by decreasing its period length. Under this condition, we found phase jump of oscillation in some simulations. Phase jump occurred only when *R* was smaller than *R*_*c*_⋅*R*_*c*_ is explained in the section below.

**Figure 2.**
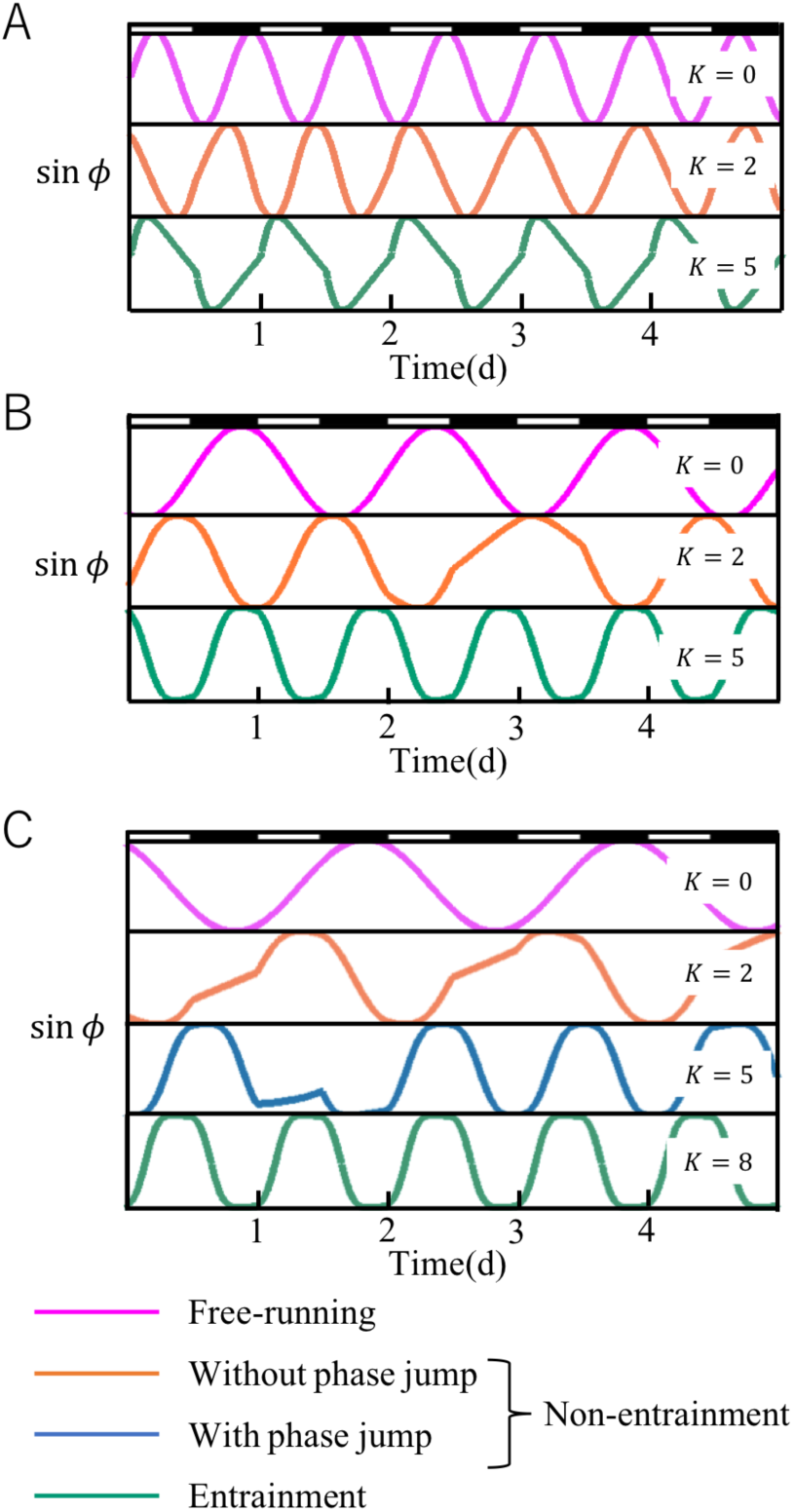
Change in oscillation behaviors with increasing *K*. Graphs in panels A, B, C show oscillations in light/dark cycles with 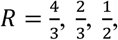, respectively. Types of behaviors are indicated by colors of lines in each graph. Open and filled rectangles denoted above graphs represent light and dark durations, respectively.

**Figure 3.**
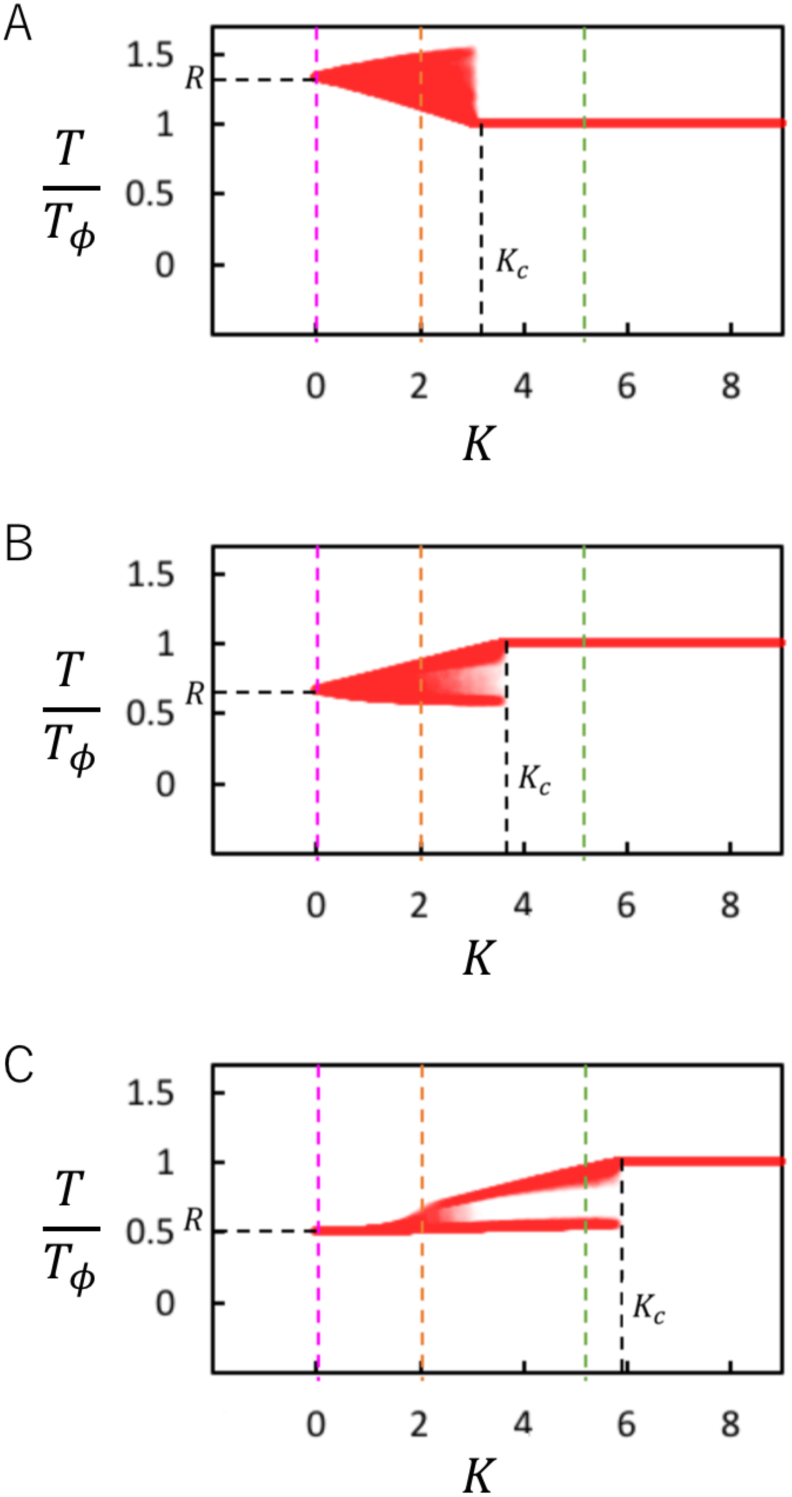
Scatter plots of *K* and the ratio of *T* to *T*_*φ*_. Here we define the length of one cycle of *φ* under light/dark cycles as *T*_*φ*_. Graphs in panels A, B, C show the scatter plots for oscillations in light/dark cycles with 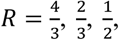 respectively. We computed *T*_*φ*_ with increasing *K* from *K* = 0 to *K* = 10 by 0.01. For each *K*, light/dark cycles were circulated for 100 times. To reduce the influence of the initial phases on the plot patterns, plots of the first five light/dark cycles in each calculation are omitted. Colored dotted lines (pink, orange, green, blue) in panels A, B, and C represent *K* values corresponding to colored wave-lines in panels A, B, C in Fig. 2, respectively.

### Critical values of *K*

We analytically calculated values of *R*_*c*_⋅*R*_*C*_ was defined as the critical value of *K* at which transition from “non-entrainment” to “entrainment” occurs. When *R* = 1, circadian oscillation is always entrained and *K*_*c*_ becomes 0. For *R* > 1, the entrainment trajectory with *K* = *K*_*c*_ includes the following points (See Appendix B):

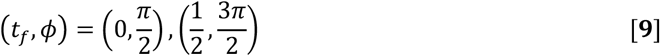

From the formula of *φ* in 0 < *K* < *ω* Eq. **7** and the boundary conditions Eq. **9**, we obtained an equation for *K*_*c*_ and *ω* as,

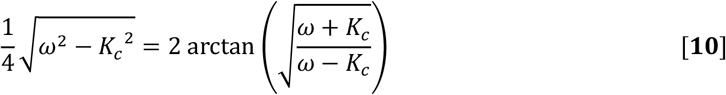

*K*_*c*_ as a solution for Eq. **10** exists when 0 < *K* < *ω* with any *R* > 1.

For *R* < 1, the entrainment trajectory with *K* = *K*_*c*_ includes the following points (See Appendix B):

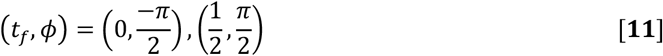

For the range of *R*_*c*_ ≤ *R* ≤ 1, *K*_*c*_ is found in 0 ≤ *K* ≤ *ω*. From the formula of *φ* in 0 ≤ *K* ≤ *ω* Eq. **7** and the boundary conditions Eq. **11**, the following equation for *K*_*c*_ and *ω* is obtained:

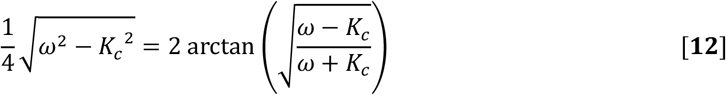

For the range of *R*_*c*_ < *R* < 1, *K*_*c*_ is found in *K* > *ω*. From the formula of *φ* in *K* > *ω* Eq. **8** and the boundary conditions Eq. **11**, the following equation for *K*_*c*_ and *ω* is obtained:

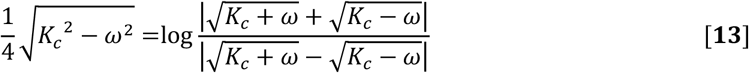

### Entrainment limits

To calculate *R*_*c*_, we analyzed the range of *R* which had *K*_*c*_ with Eq. **12**, and set a function *f*(*K*):

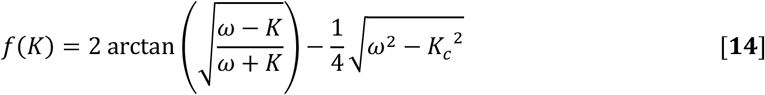

The domain of this function is defined as 0 ≤ *K* ≤ *ω*, and this function denotes a line from 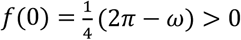.

As shown in Fig. 4, the form of *f*(*K*) was classified into two types depending on *ω*. When *ω* ≥ 4, *K*_*c*_ exists in the range of 0 ≤ *K* ≤ *ω* (Fig. 4A). This condition is derived from the following equation to find a local minimum of *f*(*K*):

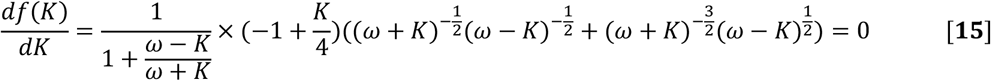

**Figure 4.**
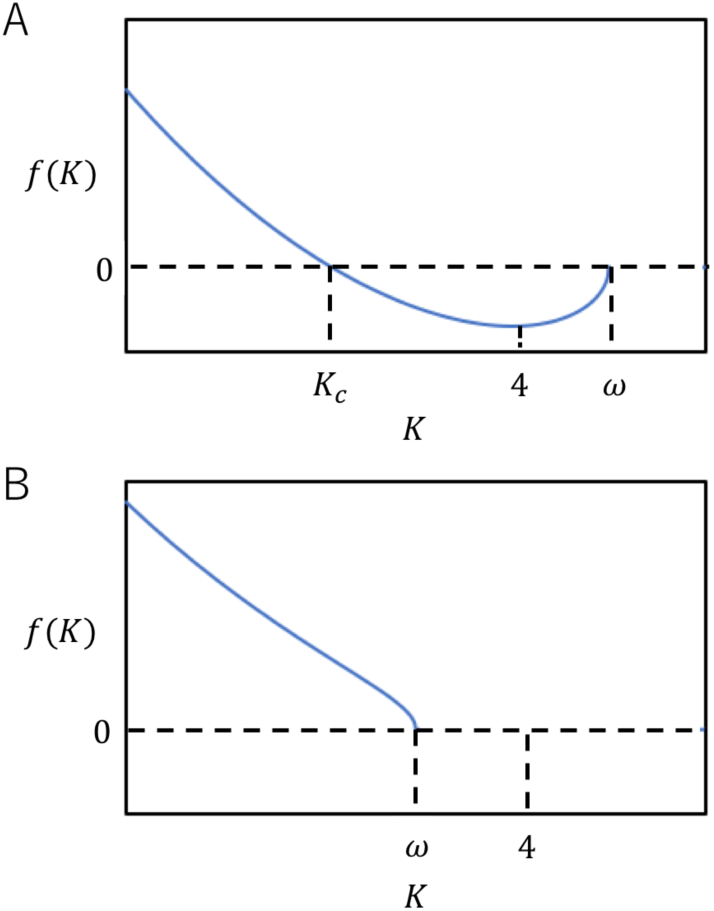
An existence condition of *K*_*c*_ defined by *f*(*K*). Representative graphs of *f*(*K*) are shown for *ω* ≥ 4 (A), *ω* < 4 (B).

This equation has a solution of *K* = 4. Therefore *ω* should be larger than or equal to 4 for existence of *K*_*c*_ in the range of 0 ≤ *K* ≤ *ω*. When *ω* < 4, *K*_*c*_ does not exist in the range of 0 ≤ *K* ≤ *ω* (Fig. 4B). Since *ω* = *2πR* is yielded from Eq. **6**, *ω* ≥ 4 means

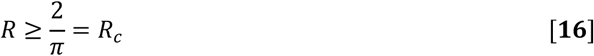

The relationship 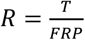 leads,

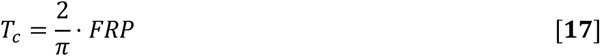

where circadian oscillation (*T* ≥ *T*_*c*_) is entrained in the range of 0 ≤ *K* ≤ *ω*. On the other hand, circadian oscillation (*T* < *T*_*c*_) is entrained in the range of *K* > *ω* with the condition of Eq. **13**.

Figure 5 shows a diagram of entrainment transition manners with *R* and *K*. Large *K* values result in entrainment of circadian rhythms under light/dark cycles (the region indicated with green). As the difference between *FRP* and *T* becomes smaller, *K*_*c*_ values become smaller. “Phase jump” (Fig. 1C) occurs in a non-entrainment region with 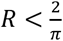 and *ω* < *K* < *K*_*c*_ (the region indicated with blue). In the other non-entrainment regions (the regions indicated with red), phase jump does not occur. Because the phase jump means a kind of reverse motion of circadian rhythm, biological relevance for these phase motions seems to be unlikely. The point 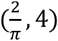 defines entrainment manners because three different entrainment manners occur around this point. We calculate the *T*_*c*_ value for the circadian rhythm with *FRP* = 24 h.

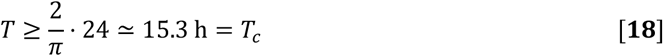

**Figure 5.**
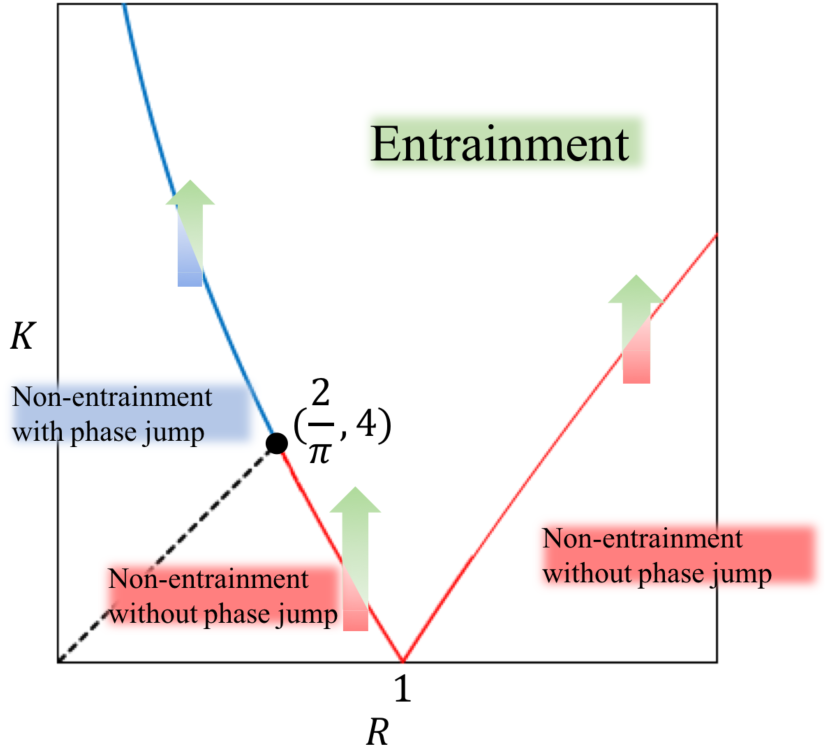
Phase diagram of the entrainment manners concerning the ratio of light/dark cycle length to free-running period (:) and the entrainment strength (*K*). The solid lines indicate the transition line for entrainment. The broken line 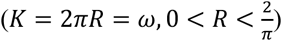 is a boundary between the manners shown in Figs. 1 and 3.

Conversely, for the entrainment in the range of 0 ≤ *K* ≤ *ω* under the condition of *T* = 24 h, period lengths of circadian oscillations are required to be shorter than the following maximum free-running period (*FRP*_*c*_)

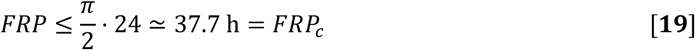

### Asymmetrical light/dark durations

As a basic setting of our modeling, we have used the light/dark cycles with the same durations of light and dark. If light/dark cycles have uneven ratios of light/dark durations, equations for *K*_*c*_ are not expressed by simple equations such as Eq. **12**. However, the range of *ω* for entrainment with 0 ≤ *K* ≤ *ω* can be rewritten (See Appendix C),

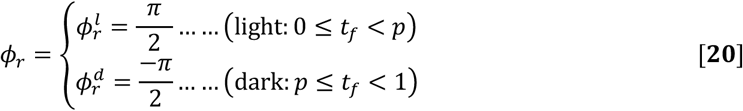

where light duration: dark duration = *p* ∶ (1 − *p*). The condition defining existence of *K*_*c*_ in 0 ≤ *K* ≤ *ω* is

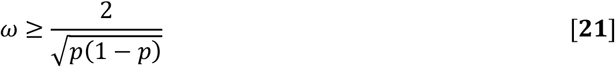

This equation is equivalent to 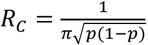 and it means

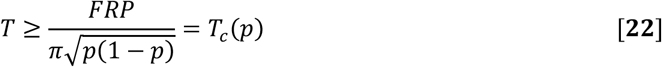

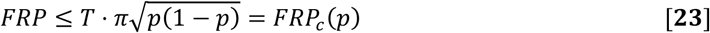

For example, *T* = 24 h and (light duration): (dark duration) = 8: 16, *FRP*_*c*_(*p*) ≃ 35.5 h. A larger deviation of *p* from 0.5 results in shorter *FRP*_*c*_(*p*). Eq. **22** indicates that this model has a limitation of the deviation of *p* for application to circadian rhythms whose *FRP*_*c*_ are longer than *T*. This limitation (*p*_*c*_) is calculated as

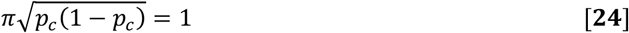

therefore,

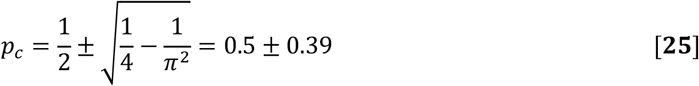

is yielded. If the ratios of light duration are shorter than 0.11 or longer than 0.89, this model is incapable of applying to circadian rhythms with *FRP* > *T*. Discrete models of phase response would be suitable.

### Comparison with previous experimental results

Capabilities of entrainment of circadian rhythms of various organisms to non-24 hour light/dark cycles (*T*-cycles) have been reported. Plants show an entrainment range between 15 and 35 hours, while mammals show much narrower ranges (10, 13). The lower limit of entrainment ranges of plants is comparable to the *T*_*c*_ value of 15.3 hours that was calculated by Eq. **18** for the circadian rhythm with *FRP* = 24 h. Circadian systems of plants showing the lower limit of approximately 15 hours are likely to have strong coupling between the light/dark signals and circadian oscillations. Circadian systems with strong coupling may contain the cellular circadian clock with a cell-autonomous light/dark signaling pathway from photoreceptors. In fact, circadian oscillation and its light input system are basically cell-autonomous in plants (11, 21). Okada et al. (2017) revealed heterogeneous entrainment behaviors between individual cellular clocks in *Lemna gibba* plants under *T*-cycles (11). In these experiments, almost all cellular rhythms of *AtCCA*1::*LUC* were entrained under *T* = 24 h, 20 h, while 54% of cellular rhythms in the same plant body were not entrained under *T* = 16 h. Under *T* = 12 h, few cellular rhythms were entrained. These heterogeneous entrainment manners appeared to reflect cell-autonomous properties of circadian systems with light inputs in plants. In contrast to plants, limited cells or tissues carry light perception systems in mammals (2). In our model, narrower entrainment ranges of mammalian circadian systems than those of plants (10) lead to much smaller *K* values. This may be caused by non-cell-autonomous properties for light inputs in mammalian circadian systems and relatively strong intercellular couplings of circadian oscillations at a tissue level.

It should be noted that the heterogeneous entrainment manners around the entrainment limit of cellular rhythms observed in plants are qualitatively represented in the present numerical results where *K* is slightly less than *K*_*c*_ in Fig. 3. Some portions of circadian behaviors both in experiments (around *T* = 16 h) and in our model (around *K*_*c*_) showed non-entrainment fluctuations with successive long and short periods (11). Furthermore, the ensemble ratios of entrainment and non-entrainment cells of *Lemna gibba* in *T* = 12 h, 16 h, 20 h, 24 h (11) seemed comparable with the theoretical calculations from our model based on experimentally-yielded *FRP* distributions of constant light (or dark) conditions (Fig. 6). *FRP*s of cellular circadian rhythms were 23.6 h ± 2.3 h under the constant light, and 30.1 h ± 2.0 h under the constant dark (11). In our model, circadian rhythms showing the light 9:; distribution are almost completely entrained under *T* = 24 h, 20 h, and partly entrained under *T* = 16 h (15%). Under *T* = 12 h, none of the cellular rhythms are entrained. Circadian rhythms showing the dark 9:; distribution are almost completely entrained under *T* = 24 h and partly entrained under *T* = 20 h (21*K*). Under *T* = 16 h, 12 h, none of the cellular rhythms are entrained. Interestingly, under each *T* value, the proportion of non-entrainment cells in experiments is a value between those calculated from our model with the light *FRP* distribution and the dark *FRP* distribution. Thus, these predictions of our continuous phase oscillator model in the conditions around *K*_*c*_ meet the entrainment manners of plant circadian rhythms.

**Figure 6.**
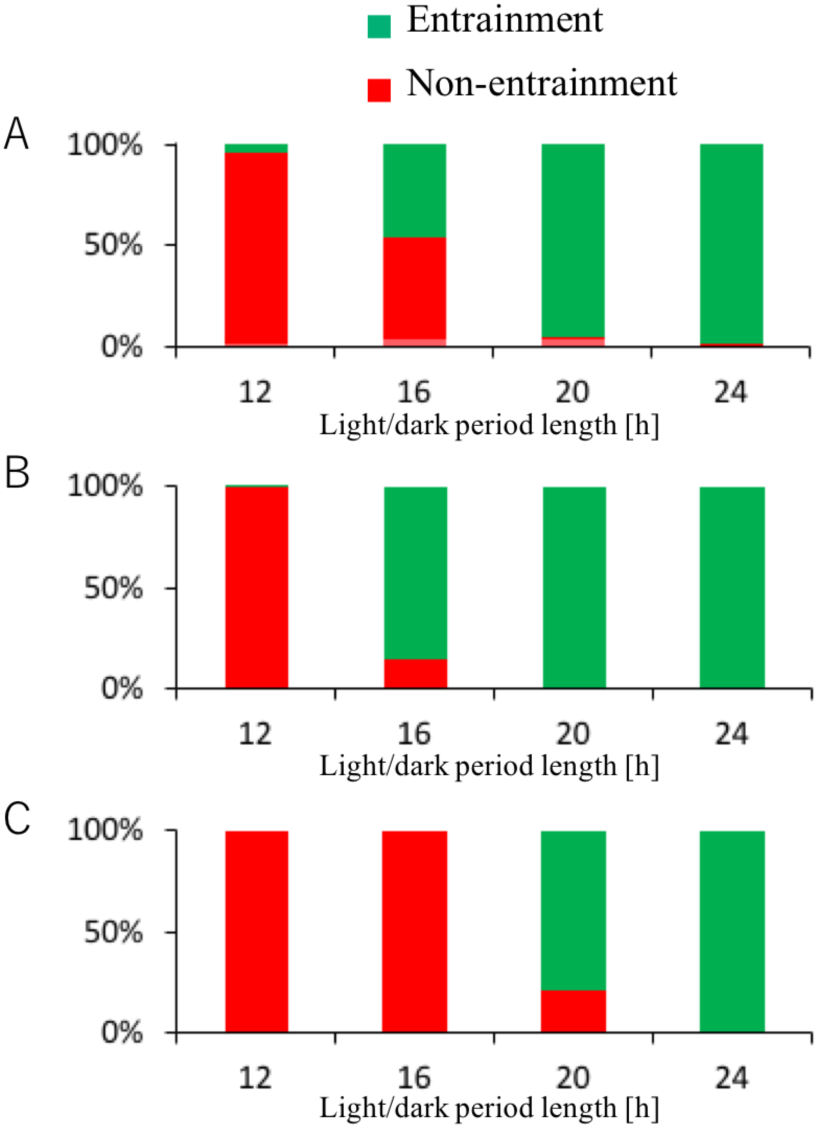
The ensemble ratios of entrainment and non-entrainment cells of *Lemna gibba* in experiments and simulation. The ratios in *T* = 12 h, 16 h, 20 h, 24 h are replotted by the experimental results (11) (A). Theoretical results by use of our model based on the experimentally-yielded FRP distributions of the plant cells measured under the constant light (B) and under the constant dark (C).

### Correspondence of our continuous model with discrete models

In most discrete models of circadian rhythms, PRC (Phase Response Curve) is used for grasping behaviors of the response function to light stimulus (1,2,7). In PRC, phases of circadian oscillations are on the x axis and phase shifts are on the y axis. Because in our model phase does not shift discretely, 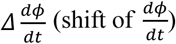 should be on the y axis instead of phase shift. Then 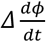 with light pulses in our model is calculated as

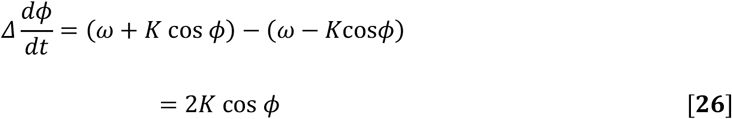

Thus in our model, phase shift by light stimulus of a short duration qualitatively follows a sinusoidal PRC. Our model may fit well with those circadian oscillations that can show a sinusoidal PRC for a light pulse. The form of phase response curve for a circadian oscillation is dependent on the strength of the light stimulus (1). The entrainment capability of a circadian oscillation to light/dark cycles is also dependent on the light intensity (22). In our model, such degrees of difference between light and dark states reflect the entrainment strength, *K*, larger degrees of difference result in larger *K* values. In this paper, we have simply denoted “light/dark cycles”, but our model can be applied to entrainment manners in any cyclic environment such as temperature cycles or feeding cycles. Entrainment manners of a circadian oscillation under a cyclic environment can be characterized by the *K*_*c*_ value that is obtained by the shorter limitation of the entrainment range of *T*-cycles. By analyzing *K*_*c*_ values, entrainment capabilities of any circadian oscillations under various cyclic environments could be deeply understood.

## Appendices

### A. Description of *φ*(*t*)

In this model, we set time scale as *T* = 1.0 day for calculation and normalize the angular velocity of circadian oscillations as 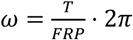. Thus, different light/dark period lengths result in different time scales of day and hour.

Case 1. *K* < *ω* and when 0 ≤ 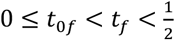

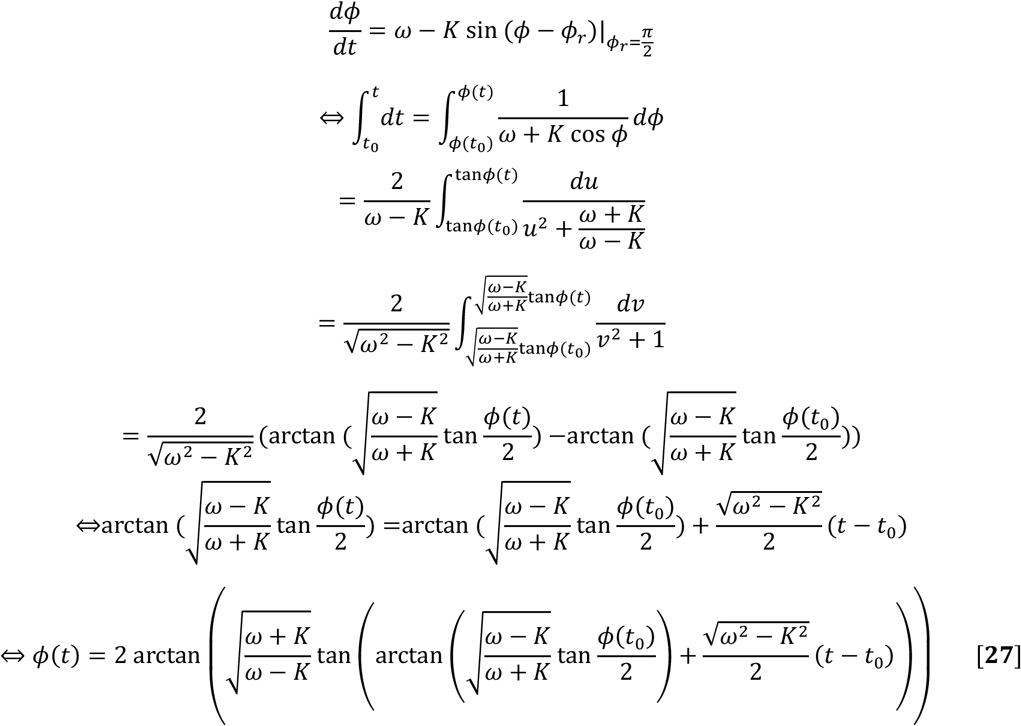

as well, when 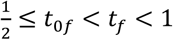

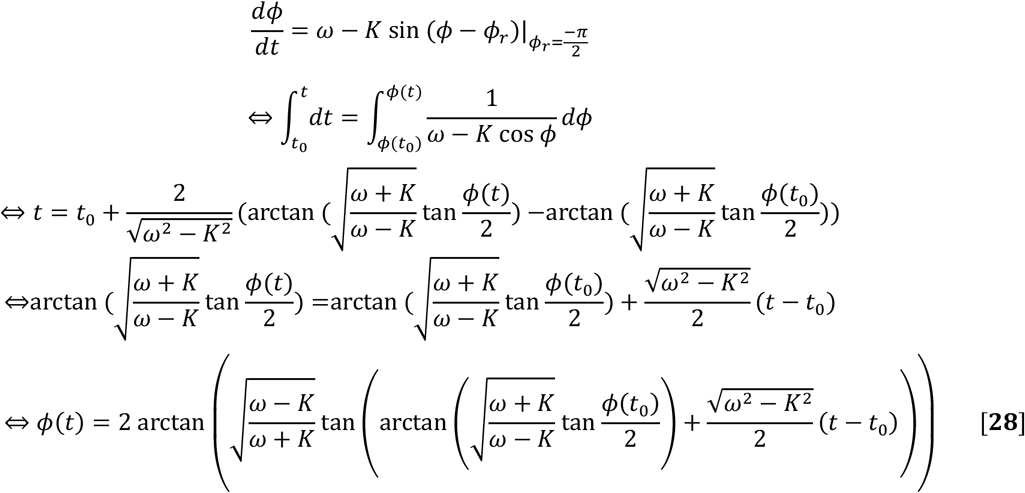

Case 2. *K* < *ω* 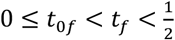

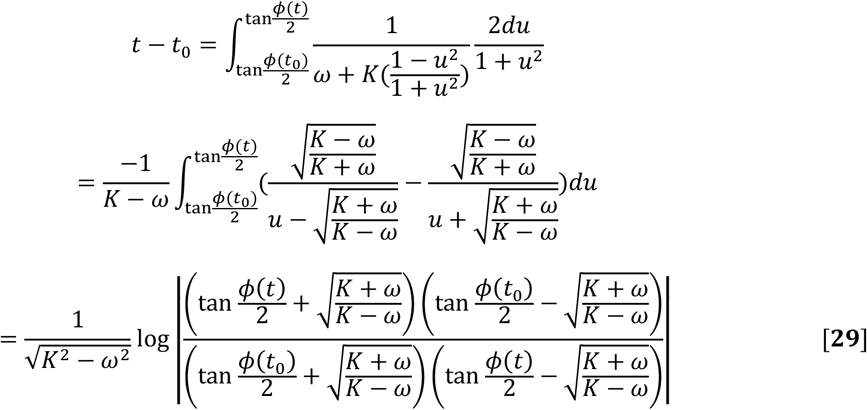

as well, when 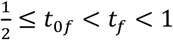

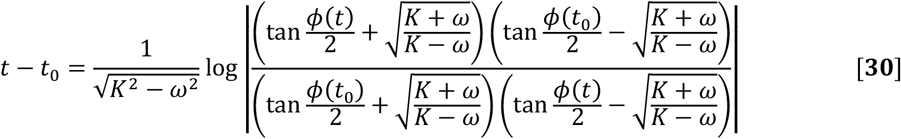

Case 3. *K* = *ω* and when 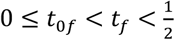

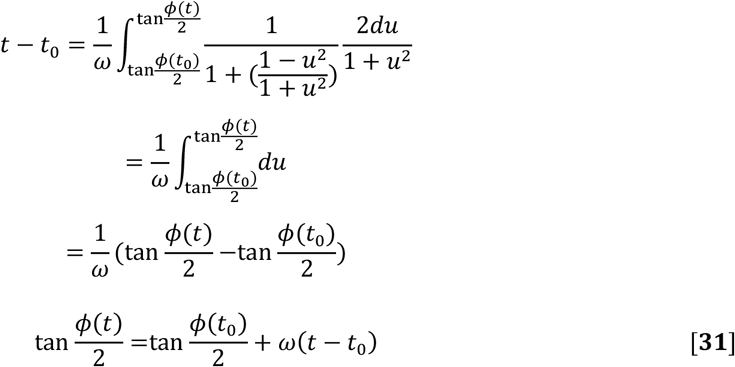

as well, when 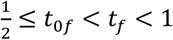

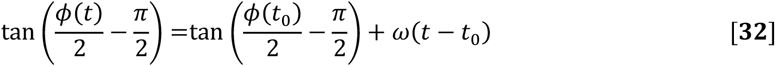

### B. Calculation of the conditions of *K*_*c*_ orbitals

Here, we describe how to calculate Eq. **9** and Eq. **11**, which are the conditions for *K* having the critical value of *K*_*c*_. Here we define the length of one cycle of *φ* under light/dark cycles as *T*_*φ*_, and also define the phase reference as *t*_*b*_(*φ*(*t*_*b*_) =0). *t*_*bf*_ is the fractional part of *t*_*b*_. From Eq. 27–32, *T*_*φ*_ is a function of *K*, *R* and *t*_*bf*_. If *FRP* < *T* and *K* < *K*_*c*_, then *T*_*φ*_ is smaller than 1 whatever value *t*_*bf*_ has. The maximum value of *T*_*φ*_ approaches 1 by increasing *K* to *K*_*c*_. When *K* reaches *K*_*c*_, the maximum value of *T*_*φ*_ becomes equal to 1. Furthermore, by denoting the *n*th *t*_*bf*_ as *t*^*n*^_*bf*_ and the *n*th *T*_*φ*_ as *T*^*n*^_*φ*_, the following equation is derived.

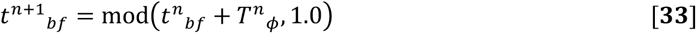

Once *T*^*n*^_*φ*_ becomes equal to 1 at *n*th,

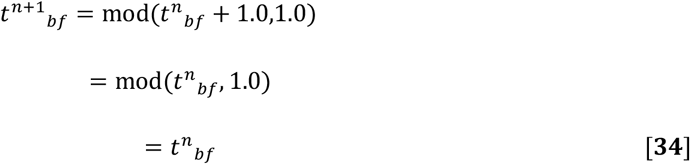

In *FRP* < *T*, Max(*T*_*φ*_) becomes equal to *T*(= 1.0), when *K* reaches a critical value (*K*_*c*_). Then, once *T*^*n*^_*φ*_ takes the Max(*T*_*φ*_) (= *T*), *T*_*φ*_ is locked at *T* from Eq. **34**. Here we define 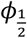 as 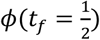 and *φ*_1_ as *φ*(*tf* = 1). At *K* = *K*_*c*_, from the symmetry of our model, *φ*_1_becomes 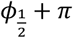. Then *T*_*φ*_ can be expressed as

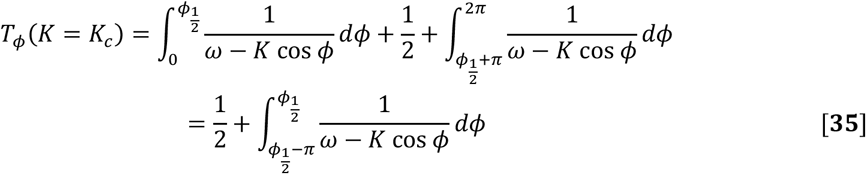

Then

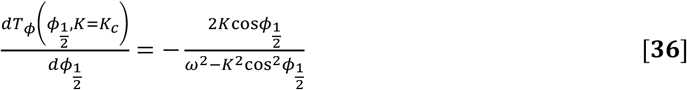

From this calculation, at *K* = *K*_*c*_

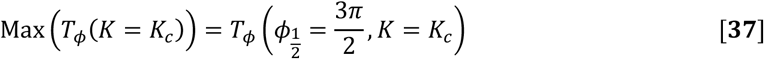

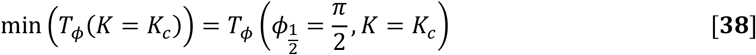

From Eq. **37**, the condition for *Max*(*T*_*φ*_) = *T* is equivalent to Eq. **9**. When *FRP* > *T*, the calculation is the same as mentioned above. From Eq. **38**, the condition for *min*(*T*_*φ*_) = *T* is equivalent to Eq. **11**.

### C. Calculation of the range of *ω* for entrainment with *K* < *ω under* light: dark ≠ 1: 1

When *K* = *ω*, from Eq. **31** and Eq. **32**, the phase of the circadian oscillation is described as

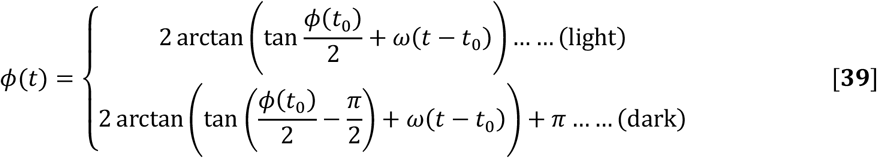

Here, we define *φ*_*p*_ as *φ*_*p*_(*t*_*f*_ = *p*) and *φ*_1_ as *φ*_1_(*t*_*f*_ = 1), respectively. From Eq. **39**,

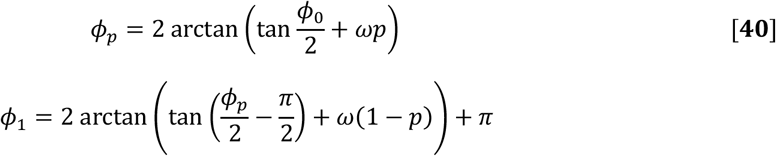

For entrainment, *φ*_1_ should be equivalent to *φ*_0_ + 2*π*.

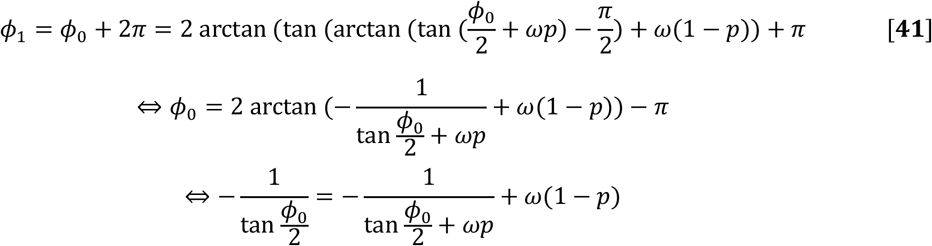

When Eq. **41** has one solution of *φ*_0_, oscillations can be entrained with *K* = *ω*. And when Eq. **41** has plural solutions, oscillations can be entrained in *K* < *ω*. Therefore, if Eq. **41** has real solutions, oscillations can be entrained in the range of 0 ≤ *K* ≤ *ω*. Then, we set 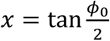, and Eq. **41** becomes

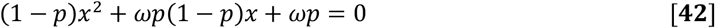

For Eq. **42** to have real solutions, its discriminant (*D*) must be larger than or equal to zero.

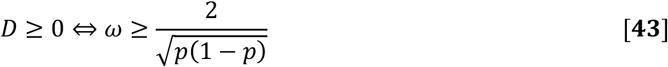

## Acknowledgements

We thank Dr. Masaaki Okada for fruitful discussions regarding the subject of this paper. We also thank the MACS program (Graduate School of Science, Kyoto University) for giving us the opportunity to undertake this research. This work was supported in part by the Japan Society for the Promotion of Science KAKENHI [grant numbers 25650098, 26707020, 17KT0022, 19H03245], and the Japan Science and Technology Agency (JST) PRESTO to T. O.

## Author contributions statement

R.T., M.I., and T.O. participated in the research design; R.T. performed the research; and R.T., M.I., and T.O. participated in writing the paper.

## References

1. C. H. Johnson, J. A. Elliott, R. Foster, Entrainment of circadian programs. Chronobiol. Int. 20, 741–774 (2003).

2. D. A. Golombek, R. E. Rosenstein, Physiology of circadian entrainment. Physiol. Rev. 90, 1063–1102 (2010).

3. J. Rémi, M. Merrow,T. Roenneberg, A circadian surface of entrainment: varying T,τ and photoperiod in *Neurospora crassa*. J. Bio. Rhythm. 25, 318–328 (2010).

4. T. Roenneberg, J. Rémi, M. Merrow, Modeling a circadian surface. J. Bio. Rhythm. 25, 340–349 (2010).

5. A. E. Granada, T. Cambras, A. D. Noguera, H. Herzel, Circadian desynchronization. Interface. Focus. 1, 153–166 (2010).

6. A. T. Winfree, Biological rhythms and the behavior of populations of coupled oscillators. J. Theor. Biol. 16, 15–42 (1967).

7. A. T. Winfree, The Geometry of Biological Time. (Springer, 2001).

8. Y. Kuramoto, “Self-entrainment of a population of coupled non-linear oscillators” in International Symposium on Mathematical Problems in Theoretical Physics. (Springer, 1975), pp. 420–422.

9. Y. Kuramoto, Chemical Oscillations, Waves, and Turbulence. (Dover Publications, 2003).

10. J. Aschoff, H. Pohl, Phase relations between a circadian rhythm and its zeitgeber within the range of entrainment. Naturwissenschaften. 65, 80–84 (1978).

11. M. Okada, T. Muranaka, S. Ito, T. Oyama, Synchrony of plant cellular circadian clocks with heterogeneous properties under light/dark cycles. Sci. Rep. 7, 317 (2017).

12. A. Graf, A. Schlereth, M. Stitt, A. M. Smith, Circadian control of carbohydrate availability for growth in *Arabidopsis* plants at night. Proc. Natl. Acad. Sci. U.S.A. 107, 9458–9463 (2010).

13. A. N. Dodd et al., Plant circadian clocks increase photosynthesis, growth, survival, and competitive advantage. Science. 309, 630–633 (2005).

14. M. A. Woelfle, Y. Ouyang, K. Phanvijhitsiri, C. H. Johnson, The adaptive value of circadian clocks: an experimental assessment in cyanobacteria. Curr. Biol. 14, 1481–1486 (2004).

15. L. C. Roden, H. R. Song, S. Jackson, K. Morris, I. A. Carre, Floral responses to photoperiod are correlated with the timing of rhythmic expression relative to dawn and dusk in *Arabidopsis*. Proc. Natl. Acad. Sci. U.S.A. 99, 13313–13318 (2002).

16. A. N. Dodd, N. Dalchau, M. J. Gardner, S. J. Baek, A. A. Webb, The circadian clock has transient plasticity of period and is required for timing of nocturnal processes in Arabidopsis. New. Phytol. 201, 168–179 (2014).

17. A. Erzberger, G. Hampp, A. E. Granada, U. Albrecht, H. Herzel, Genetic redundancy strengthens the circadian clock leading to a narrow entrainment range. J. Roy. Soc. Interface. 10, 20130221 (2013).

18. H. G. McWatters, R. M. Bastow, A. Hall, A. J. Millar, The *ELF3 zeitnehmer* regulates light signalling to the circadian clock. Nature. 408, 716–720 (2000).

19. G. B. Ermentrout, J. Rinzel, Beyond a pacemaker’s entrainment limit: phase walk through. Am. J. Physiol. Regulatory Integrative Comp. Physiol. 246, R102–R106 (1984).

20. S. H. Strogatz, Nonlinear Dynamics and Chaos: with Applications to Physics, Biology, Chemistry, and Engineering. (Westview Press, 2014).

21. T. Muranaka, T. Oyama, Heterogeneity of cellular circadian clocks in intact plants and its correction under light-dark cycles. Sci. Adv. 2, e1600500 (2016).

22. C. Gronfier, K. P. Wright, R. E. Kronauer, C. A. Czeisler, Entrainment of the human circadian pacemaker to longer-than-24-h days. Proc. Natl. Acad. Sci. U.S.A. 104, 9081–9086 (2007).

